# Incorporation of data from multiple hypervariable regions when analyzing bacterial 16S rRNA sequencing data

**DOI:** 10.1101/2021.06.17.448728

**Authors:** Carli B. Jones, James R. White, Sarah E. Ernst, Karen S. Sfanos, Lauren B. Peiffer

## Abstract

Short read 16S rRNA amplicon sequencing is a common technique used in microbiome research. However, inaccuracies in estimated bacterial community composition can occur due to amplification bias of the targeted hypervariable region. A potential solution is to sequence and assess multiple hypervariable regions in tandem, yet there is currently no consensus as to the appropriate method for analyzing this data. Additionally, there are many sequence analysis resources for data produced from the Illumina platform, but fewer open-source options available for data from the Ion Torrent platform. Herein, we present an analysis pipeline using an open-source analysis platform that integrates data from multiple hypervariable regions and is compatible with data produced from the Ion Torrent platform. We used the ThermoFisher Ion 16S™ Metagenomics Kit and a mock community of 20 bacterial strains to assess taxonomic classification of amplicons from 6 separate hypervariable regions (V2, V3, V4, V6-7, V8, V9) using our analysis pipeline. We report that different hypervariable regions have different specificities for taxonomic classification, which also had implications for global level analyses such as alpha and beta diversity. Finally, we utilize a generalized linear modeling approach to statistically integrate the results from multiple hypervariable regions and apply this methodology to data from a small clinical cohort. We conclude that scrutinizing sequencing results separately by hypervariable region provides a more granular view of the taxonomic classification achieved by each primer set as well as the concordance of results across hypervariable regions. However, the data across all hypervariable regions can be combined using generalized linear models to statistically evaluate overall differences in community structure and relatedness among sample groups.

## Introduction

Next generation sequencing of microbial DNA has become an important tool used for determining relationships between human-associated microbial populations and various diseases. Most studies in this realm rely on either shotgun metagenomic sequencing or 16S ribosomal RNA (rRNA) amplicon sequencing. Shotgun metagenomic sequencing involves sequencing random fragments of sample DNA which contains a mixture of bacterial DNA, as well as host and other microbial and environmental DNA (1). This method allows for taxonomic profiling, metabolic function profiling, and antibiotic resistance gene profiling; however, it is generally more expensive than amplicon sequencing, and requires a large amount of input DNA and the availability of reference genome sequences. Bacterial 16S rRNA amplicon sequencing employs PCR amplification of specific hypervariable regions within the gene, followed by deep sequencing (2). This method is generally a quicker, cheaper alternative to shotgun metagenomics; however, it only identifies bacteria and the typical strategy only sequences a specific fragment of the bacterial 16S rRNA gene (3). The 16S rRNA gene is comprised of 9 hypervariable regions (V1-V9), and most primers used for next generation sequencing only target one to two hypervariable regions at a time. Multiple studies have shown that different regions vary in their taxonomic utility due to a combination of primer bias, differential hypervariable region sequence length, and hypervariable region sequence uniqueness across bacterial taxa (4–8). An ideal solution would be to sequence the entire 16S rRNA gene, however this requires long-read sequencing that is very time and resource intensive. Therefore, a potential alternative would be to perform 16S rRNA amplicon sequencing on multiple regions and incorporate information from as many hypervariable regions as possible into downstream data analysis.

The Ion 16S™ Metagenomics Kit (Life Technologies) utilizes primers to seven different hypervariable regions: V2, V3, V4, V6-7, V8, and V9. This is an attractive approach because it yields more sequence information across the 16S rRNA gene overall. However, there is currently little consensus as to how to properly analyze information from multiple hypervariable regions and obtain overall results. Current analysis pipelines for Ion Torrent data include the Ion Reporter Software offered by ThermoFisher, and an alternative method using open access tools developed by Barb *et al.* (5). Both pipelines offer methods for taxonomic identification; however, they do not address the question of how to appropriately integrate data from multiple hypervariable regions in downstream analyses. Recently, Fuks *et al.* (9) and Debelius *et al.* (10) developed methods to computationally combine data from multiple hypervariable regions to provide a joint estimate of the microbial community composition. To date, however, there is no generally agreed upon approach for combining sequences from multiple hypervariable regions for downstream analyses, especially for less commonly used 16S rRNA sequencing platforms such as Ion Torrent.

Herein, we developed an analysis pipeline that analyzes data from each hypervariable region separately, allowing for systematic comparison of taxonomic classification by hypervariable region. We demonstrate our results from analyzing a mock community of bacterial DNA where we determine how each hypervariable region differs in its utility to provide information on taxonomic classifications, alpha diversity, and beta diversity. We report that certain taxa are only identified by particular V regions, supporting our hypothesis that there is a benefit to incorporating multiple primer sets into sequencing strategies. Furthermore, we discuss different options for downstream analysis and statistics, and propose a generalized linear model (GLM) to statistically combine the results from multiple hypervariable regions. Finally, we demonstrate the utility of our approach in the analysis of clinical samples.

## Materials and methods

### Clinical sample collection and mock community

All specimens were studied under an Institutional Review Board (IRB) approved protocol with written informed consent. Three adult males self-collected two rectal swab samples with sterile flocked swabs (Copan Diagnostics, Murrieta, CA). DNA was extracted from one of the swabs from each individual immediately after collection (referred to as RS1). The other swabs (referred to as RS2) were frozen at −80°C for six days before DNA extraction.

The 20 Strain Even Mix Genomic Material was obtained from American Type Culture Collection (ATCC, Cat. No. MSA-1002, Manassas, VA). The strain composition of the mock community is given in Table 1. The mock community was sequenced with five technical replicates.

**Table 1.** Contents of mock community.

| Species | 16S<br>Copies<sup>a</sup> | Genus | Family |
| --- | --- | --- | --- |
| *Acinetobacter baumannii* | 6 | *Acinetobacter* | *Moraxellaceae* |
| *Actinomyces odontolyticus* | 2 | *Actinomyces* | *Actinomycetaceae* |
| *Bacillus cereus* | 12 | *Bacillus* | *Bacillaceae* |
| *Bacteroides vulgatus* | 7 | *Bacteroides* | *Bacteroidaceae* |
| *Bifidobacterium adolescentis* | 5 | *Bifidobacterium* | *Bifidobacteriaceae* |
| *Clostridium beijerinckii* | 14 | *Clostridium* | *Clostridiaceae* |
| *Cutibacterium acnes* | 4 | *Cutibacterium* | *Propionibacteriaceae* |
| *Deinococcus radiodurans* | 7 | *Deinococcus* | *Deinococcaceae* |
| *Enterococcus faecalis* | 4 | *Enterococcus* | *Enterococcaceae* |
| *Escherichia coli* | 7 | *Escherichia* | *Enterobacteriaceae* |
| *Helicobacter pylori* | 2 | *Helicobacter* | *Helicobacteraceae* |
| *Lactobacillus gasseri* | 6 | *Lactobacillus* | *Lactobacillaceae* |
| *Neisseria meningitidis* | 4 | *Neisseria* | *Neisseriaceae* |
| *Porphyromonas gingivalis* | 4 | *Porphyromonas* | *Porphyromonadaceae* |
| *Pseudomonas aeruginosa* | 4 | *Pseudomonas* | *Pseudomonadaceae* |
| *Rhodobacter sphaeroides* | 3 | *Rhodobacter* | *Rhodobacteraceae* |
| *Staphylococcus aureus* | 6 | *Staphylococcus* | *Staphylococcaceae* |
| *Staphylococcus epidermidis* | 5 | *Staphylococcus* | *Staphylococcaceae* |
| *Streptococcus agalactiae* | 7 | *Streptococcus* | *Streptococcaceae* |
| *Streptococcus mutans* | 5 | *Streptococcus* | *Streptococcaceae* |

<sup>a</sup>Number of copies of 16S rRNA contained in the bacterial genome of the indicated species.

### DNA extraction

The DNA extraction protocol was adapted from our previously published protocol (11). Briefly, rectal swab fecal material was resuspended in 500 μl of 1X phosphate buffered saline (PBS). Samples were then digested in a cocktail of lysozyme (10 mg/ml, Sigma-Aldrich, St. Louis, MO) and mutanolysin (25 KU/ml, Sigma-Aldrich) for 1 hour at 37 °C. The contents of the tubes were then transferred into FastPrep Lysing Matrix B tubes (MP Biomedicals, Santa Ana, CA). Next, 20% SDS (Sigma-Aldrich) and phenol:chloroform:isoamyl alcohol (25:24:1, ThermoFisher Scientific, Waltham, MA) were added and samples were homogenized by bead beating in an MP FastPrep-24 at 6m/s for a total of 60 seconds. DNA was precipitated and resuspended in a final volume of 50 μl of DNA-free water (Molzym, Bremen, Germany).

### Library preparation

Concentration of DNA from the mock microbial community (Table 1) and rectal swabs was measured using a Qubit dsDNA HS (high sensitivity) kit (Life Technologies, Carlsbad, CA). Libraries were prepared using the Ion 16S™ Metagenomics Kit (Cat. No. A26216, ThermoFisher Scientific). Briefly, 10 ng of DNA was mixed with 15 μl of Environmental Master Mix. 3 μl of each 16S Primer Set (10X) was added to each tube, one sample set with primers for V2-4-8 (Pool 1) and the other with primers for V3-6,7-9 (Pool 2). Samples were placed in a thermocycler with the following thermal conditions: 95°C for 10 minutes, 25 cycles of 95°C for 30s, 58°C for 30s, 72°C for 30s, then 72°C for 7min. Amplification products were purified using AMPure XP beads (Cat. No. A63881, Beckman Coulter, Pasadena, CA) and eluted in nuclease free water. Concentrations of amplification products from Pool 1 and Pool 2 were measured using a Bioanalyzer and the High Sensitivity DNA Kit (Agilent Technologies, Santa Clara, CA), and the two pools were combined for a total of 100 ng of DNA (50ng from each pool).

Next, 20 μl of 5X End Repair Buffer and 1 μl of End Repair Enzyme were added to each sample, and then incubated for 20 min at room temperature. Pooled amplicons were then purified again using AMPure XP beads and eluted in Low TE buffer. Ligation and nick repair were performed using 10X Ligase Buffer, Ion P1 Adaptor, Ion Xpress Barcodes, dNTP Mix, DNA Ligase, Nick Repair Polymerase, nuclease-free water, and sample DNA with the following thermal conditions: 25°C for 15min, 72°C for 5min. Adapter-ligated and nick-repaired DNA was then purified using AMPure XP beads and eluted in Low TE buffer.

The library was then amplified using the Ion Plus Fragment Library Kit (Cat. No. 4471252, ThermoFisher Scientific) with the following thermal conditions: 95°C for 5min, 7 cycles of 95°C for 15s, 58°C for 15s, 70°C for 1min, and then 70°C for 1min. The amplified library was then purified using AMPure XP beads and eluted in Low TE buffer. Library concentrations were measured using a Bioanalyzer and the High Sensitivity DNA Kit. Libraries were then diluted down to 26 pM and pooled, yielding a 26 pM solution.

### Sequencing

Libraries were prepared for sequencing using oil amplification to template the libraries onto beads and loaded onto chips using the Ion Chef Instrument and the Ion 520™ & Ion 530™ Kit – Chef (ThermoFisher Scientific). Chips were then loaded onto the Ion GeneStudio S5 System along with Ion S5 Sequencing Kit reagents (ThermoFisher Scientific) and sequenced. Samples in this study were sequenced across three separate sequencing runs on Ion 520 and Ion 530 chips using 400bp sequencing kits. Sequences were demultiplexed by sample using the S5 device software, and then separated per hypervariable region by ThermoFisher prior to downstream analysis.

### Database curation

When using SILVA (12) or Greengenes (13) as our reference database, approximately 50% of reference sequences were unclassified. Therefore, we utilized a refined version of the SILVA (v.123) database for taxonomic assignment with unclassified and distantly related sequences were removed. The refined database *(sfanos-db-4.0)* contains approximately 15,000 named species and includes species that are of particular interest for future clinical gastrointestinal (GI) microbiome studies.

### *In silico* taxonomic validation of curated database

To determine how well our curated database could assign taxonomy to sequences and how much the assignment differs by hypervariable region, we created a system of *in silico* taxonomic validation using a human gut microbiome culture collection (14). First, we separated the sequences in the culture collection by hypervariable region. To do this, we ran the sequences from the culture collection through NCBI BLAST against the ATCC mock community sequences that had already been split by hypervariable region. This method allowed us to break down the culture collection sequences into their different hypervariable regions and simulate more complex clinical data. A 1% noise rate was included in the simulated sequences to mimic typical evolutionary variation in species as well as sequencing error. We then ran taxonomic classification of the sequences from the culture collection using our curated database, with a threshold of 97% sequence identity. A confidence score was assigned to each classification by VSEARCH. Results were categorized into true positives, false positives, and false negatives based on whether they were in the culture collection or not.

### Data processing

The ThermoFisher Bioinformatics team divided each fastq file by hypervariable region using proprietary primer sequences to produce 6 separate fastq files per sample (V2, V3, V4, V6-7, V8, and V9), with primer sequences removed and all reads oriented in the forward direction. Manifest files were then created for each hypervariable region and each sequencing run. Fastq files were imported into QIIME2 format via qiime tools import in SingleEndFastqManifestPhred33V2 format (15). QIIME2 v 2020.6 was used to perform denoising, Operational Taxonomic Unit (OTU) clustering, taxonomic classification, phylogenetic tree construction, and alpha and beta diversity.

DADA2 was used to denoise data, using the denoise-pyro plugin and parameters of 0bp for trimming and truncation (16). A separate DADA2 run was performed for each hypervariable region and each sequencing run. Denoising statistics were then exported and summarized via the B02-summarize-qc.pl Perl script. From this summary, we determined that all samples in all hypervariable regions had a minimum of 10,000 reads which passed the filter in the DADA2 step. Good’s coverage was performed at a depth of 10,000 reads for each hypervariable region and at least 99% coverage was achieved for all regions (17). Thus, we decided that 10,000 reads was an acceptable sampling depth. DADA2 feature tables and representative sequence files were then merged across sequencing runs so that there was only one feature table and representative sequence file per hypervariable region.

Open-reference OTU clustering was then performed using QIIME2 plugin vsearch cluster-features-open-reference (18). A threshold of 99% identity was used, and sequences were clustered against reference sequences from the curated sfanos_db_v4.0.fastq.qza database.

### Alpha and beta diversity analysis

A phylogenetic tree was constructed for each hypervariable region using the representative sequences file generated from open-reference OTU clustering via the qiime phylogeny align-to-tree-mafft-fasttree plugin (19–21), Shannon diversity (22), evenness, and observed OTUs. Distance matrices were exported for Jaccard (23), Bray-Curtis (24), weighted (25), and unweighted (26) UNIFRAC distances. Distance matrices were created from filtered taxonomic abundance results comparing region to region. Visualizations of alpha diversity metrics as well as principal coordinates analysis (PCoA) plots were created using R in order to determine region to region differences.

### Taxonomic classification

Taxonomic classification was performed using classify-consensus-vsearch using the curated sfanos_db_v4.0 reference reads and reference taxonomy with 99% identity. The output .qza file was then exported in order to obtain the taxonomy.tsv file. This file and the feature-table.biom file were used in a Perl script designed to summarize the taxonomic information into feature-table-with-taxonomy.txt. Heatmaps were created in R using the pheatmap package and taxa-normalize-pct-per-region.txt file.

### Contaminant filtering

Contaminant sequences were filtered out from the ATCC sample data. Any taxa that were detected in only one of the five technical replicates, detected at less than 0.1% abundance, or both, was considered a contaminant. Filtering was performed on the feature table that was created after open reference OTU clustering using qiime taxa filter-table. Contaminants are listed in S1 Table.

### Generalized linear modeling

We used the generalized linear model function in Base R to evaluate statistical differences in alpha diversity and individual taxonomic abundance between fresh versus frozen samples in the clinical cohort. The GLM per feature took the following structure: *log10(feature) ~ fresh/frozen status + specimen ID + hypervariable region.* Regions V8 and V9 were excluded from GLM analysis, and Region V2 was used as the null factor level. The fresh/frozen status of samples was compared, with fresh as baseline factor level set as zero and frozen set as one. The input of “feature” was either an alpha diversity value (Shannon, Evenness, Observed OTUs or Faith’s Phylogenetic Diversity), or taxonomic abundance of a feature at a specific taxonomic level. Input feature values were log transformed in order to increase stability of values from person to person when performing statistics. The GLM p-value was obtained by comparing the GLM factor level coefficient to the null hypothesis of zero, which was done via a Wald Test.

### Data and code availability

All sequences were submitted to the NCBI Sequence Read Archive (SRA) under Bioproject ID PRJNA738491. Sequences will be released upon publication in a peer-reviewed journal. All codes used can be found on the public GitHub repository it-workflow (http://github.com/Sfanos-Lab-Microbiome-Projects/it-workflow/), which will be made public upon publication in a peer-reviewed journal.

## Results

### *In silico* validation of taxonomic classification accuracy of curated database

We curated a database specific for our needs during our pipeline development using SILVA as the base. We then validated the classification accuracy of this database by performing an *in silico* analysis where we ran known sequences from a human gut microbiome culture collection (14) through our classification scheme and recorded whether the sequences were classified correctly (true positive), incorrectly (false positive), or not identified (false negative). Results were compiled by unique sequence identifier in a pivot table which noted the corresponding hypervariable region, true taxonomic classification at the genus and species levels, taxonomic classification by VSEARCH using our curated database, VSEARCH confidence score, and classification of assignment by VSEARCH at both the genus and species level as either True Positive (TP), False Positive (FP), or False Negative (FN) (S1 File).

Sequence assignment counts were converted to percent by adding up the total number of sequences that were assigned as TP, FP, or FN for each V region, dividing by the total number of sequences for that region, and multiplying by 100. Taxonomic classification across all V regions was more accurate at the genus level than the species level, with a combined total of 85.08% TP, 6.95% FP, and 7.97% FN verses 65.96% TP, 10.31% FP, and 23.74% FN, respectively. Results were not evenly distributed across V regions. V3 had the highest accuracy at the genus level in all three metrics whereas V2, V4, and V6-7 performed similarly to each other and slightly less accurately than V3. V8 and V9 had lower TP and higher FN and FP than all other V regions (Table 2). At the species level, regions V2 and V3 had the highest accuracy, with low FP and FN counts and highest TP counts. Regions V8 and V9 had much lower accuracy with the highest FP and FN counts and lowest TP counts. Regions V4 and V6-7 performed in between (Table 2).

**Table 2.** Results of *in silico* analysis.

| Genus Level |  |  |  |
| --- | --- | --- | --- |
| Region | False Neg | False Pos | True Pos |
| V2 | 4.85% | 4.62% | 90.53% |
| V3 | 2.82% | 1.99% | 95.19% |
| V4 | 6.22% | 6.71% | 87.07% |
| V6-7 | 5.22% | 5.79% | 88.98% |
| V8 | 6.96% | 14.50% | 78.54% |
| V9 | 21.96% | 8.14% | 69.91% |
| Species Level |  |  |  |
| Region | False Neg | False Pos | True Pos |
| V2 | 13.48% | 8.58% | 77.94% |
| V3 | 16.24% | 6.10% | 77.66% |
| V4 | 25.84% | 8.12% | 66.04% |
| V6-7 | 16.58% | 10.03% | 73.39% |
| V8 | 23.83% | 17.46% | 58.71% |
| V9 | 46.83% | 11.56% | 41.61% |

Ability of curated database to accurately classify sequences from each hypervariable region from known taxa at the genus and species level in an *in silico* analysis.

### Mock community

In order to test our analysis pipeline, we prepared libraries and sequenced DNA from a mock microbial community (20 Strain Even Mix Genomic Material, ATCC® MSA-1002™). Five independent replicate library preparations of the mock community were sequenced. We filtered out low-level contaminants (S1 Table) prior to performing community alpha and beta diversity and taxonomic abundance analyses (see Methods).

### Alpha diversity

We analyzed 4 different alpha diversity metrics, 2 measures of evenness (Evenness and Shannon Diversity), and 2 measures of richness (Faith’s Phylogenetic Diversity and Observed-OTUs) (Fig 1). V9 has significantly decreased alpha diversity compared to all regions across all metrics (S2 File). V8 also has significantly decreased Shannon diversity, Evenness, and Faith’s phylogenetic diversity compared to other regions excluding V9, with two exceptions being that Evenness is not significantly decreased in V8 compared to that of V6-7 and Faith’s PD is not significantly decreased in V8 compared to V4 (S2 File).

**Fig 1.**
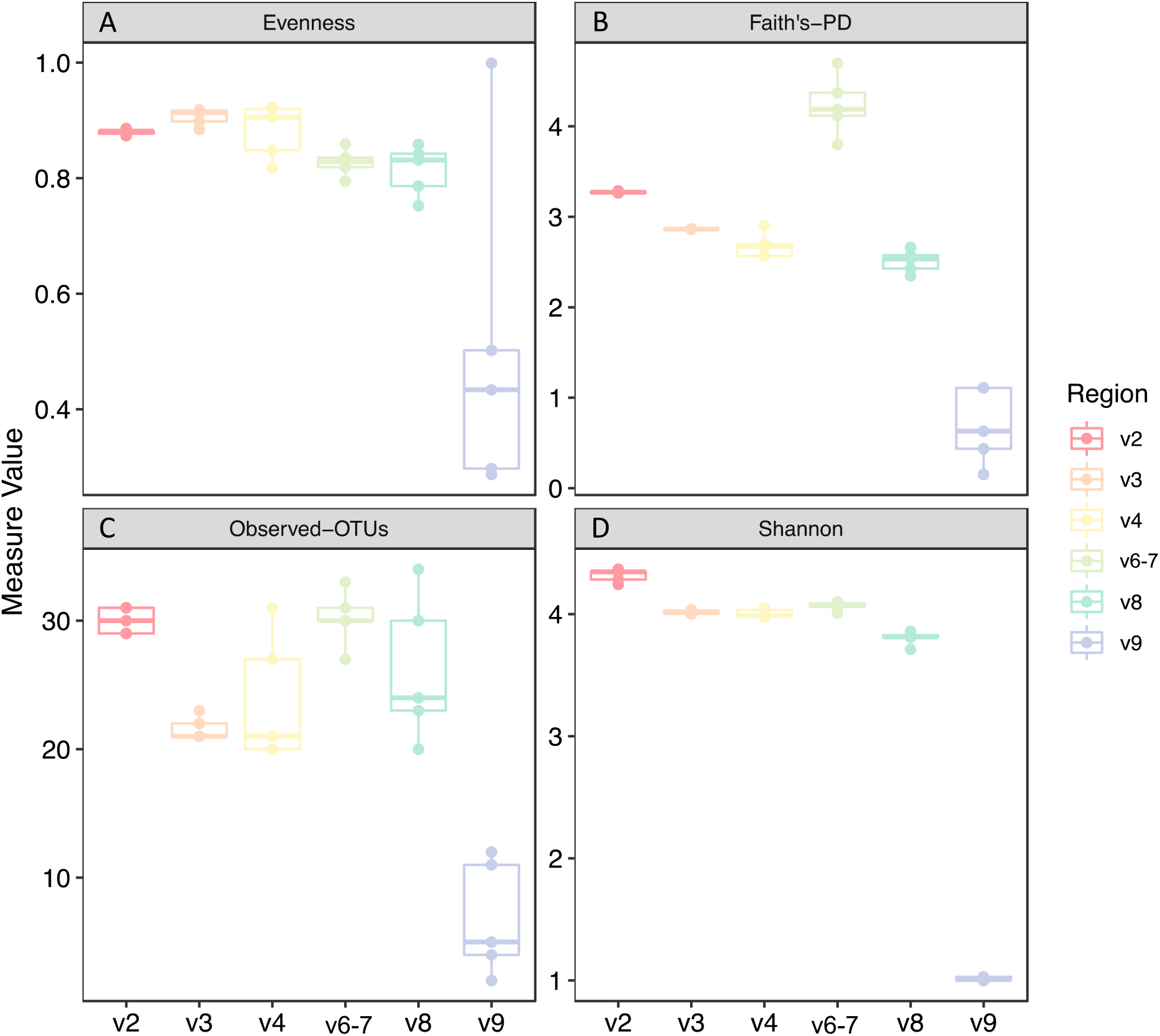
Alpha diversity analyses of mock community technical replicates by hypervariable region. Evenness (A), Faith’s phylogenetic diversity (B), Observed Operational Taxonomic Units (OTUs) (C), Shannon diversity (D).

### Beta diversity

To compare beta diversity between hypervariable regions and circumvent the issue that OTUs would be region-specific, we used taxonomic results from each hypervariable region to create aggregated distance matrices. We assessed six different beta diversity metrics: Canberra, Bray-Curtis, Jaccard, Euclidean, Gower, and Kulczynski. Fig 2 shows a PCoA plot based on the Canberra distance matrix. The V2, V3, V4, and V6-7 hypervariable regions clustered together, whereas V8 and V9 were distantly separated. This pattern was observed by other beta diversity metrics, with V6-7 sometimes also segregating slightly from V2, V3, and V4 which were largely clustered together (S1 Fig).

**Fig 2.**
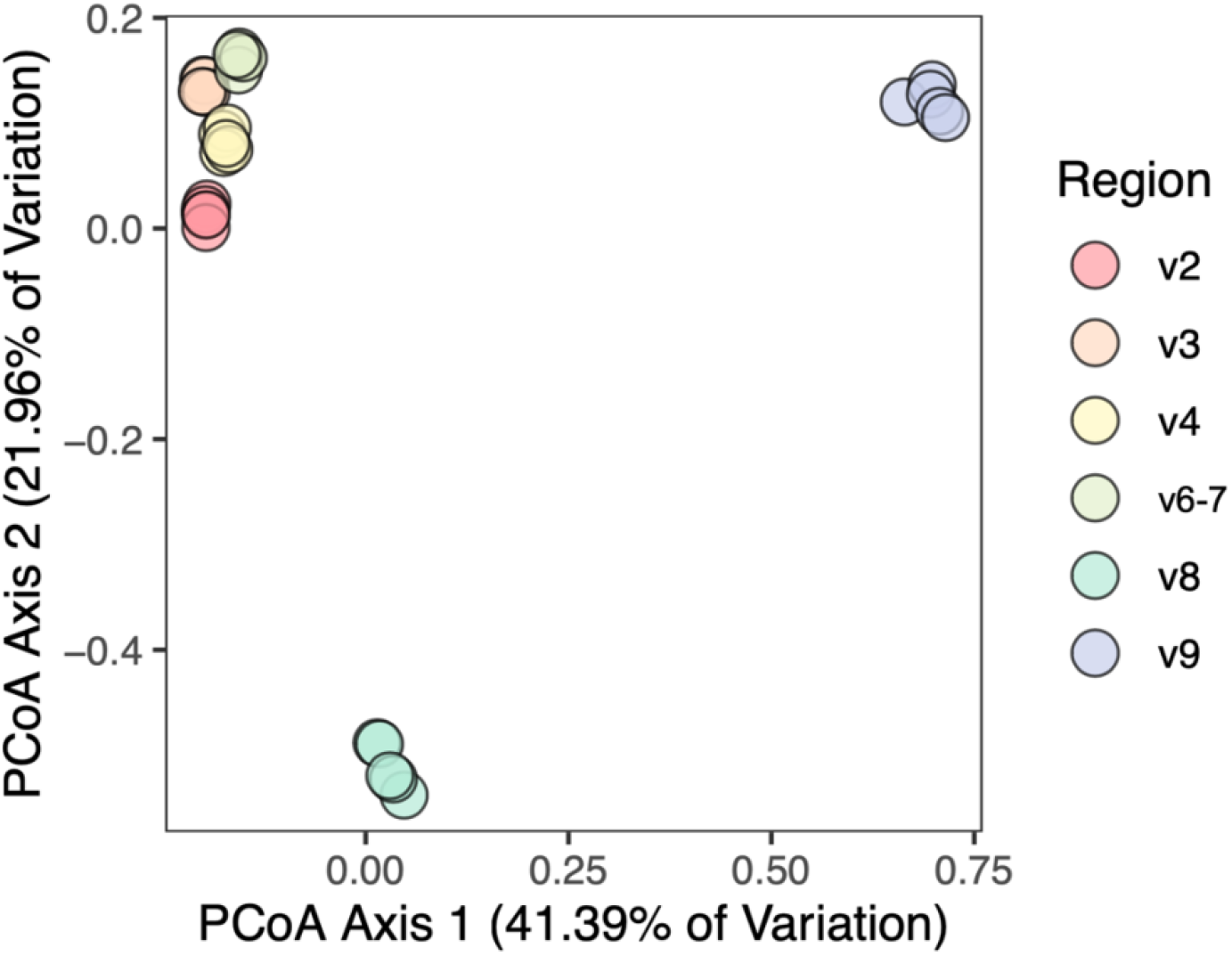
Principal coordinates analysis of mock community samples using Canberra distance matrix.

### Taxonomic classification and abundance

The percent abundance of the identified organisms after taxonomic classification is given in S3 File. The majority of species were identified by taxonomic classification of the sequences covering each hypervariable region, with the exception of V9 that only positively identified *Escherichia coli* and *Acinetobacter baumannii. Clostridium beijerinckii* was the most difficult organism to speciate and was only correctly classified in V6-7 amplicons. The results with hypervariable regions V2, V3, and V4 only identified *Clostridium beijerinckii* at the genus level, V8 mis-classified it as *Clostridium butyricum*, and V9 did not identify any Clostridial organisms (S3 File). Aside from *C. beijerinckii,* species misclassification varied by hypervariable region.

We next compared observed versus expected percent abundance by hypervariable region. To determine the expected percent abundance, we divided the number of 16S rRNA copies present in each bacterium (Table 1) and divided by the total number of 16S rRNA copies in the 20 Strain Even Mix. Taxonomic bar plots demonstrate the percent abundance of each taxa by hypervariable region compared to expected (Fig 3). V2 most closely approximated the overall distribution of species compared to expected and correctly assigned the most species from the mock community (19/20). V3 (17/20), V6-7 (17/20), and V4 (16/20) followed closely behind, whereas V8 assigned 15/20, and V9 was only able to identify two species (2/20) (Table 3).

**Fig 3.**
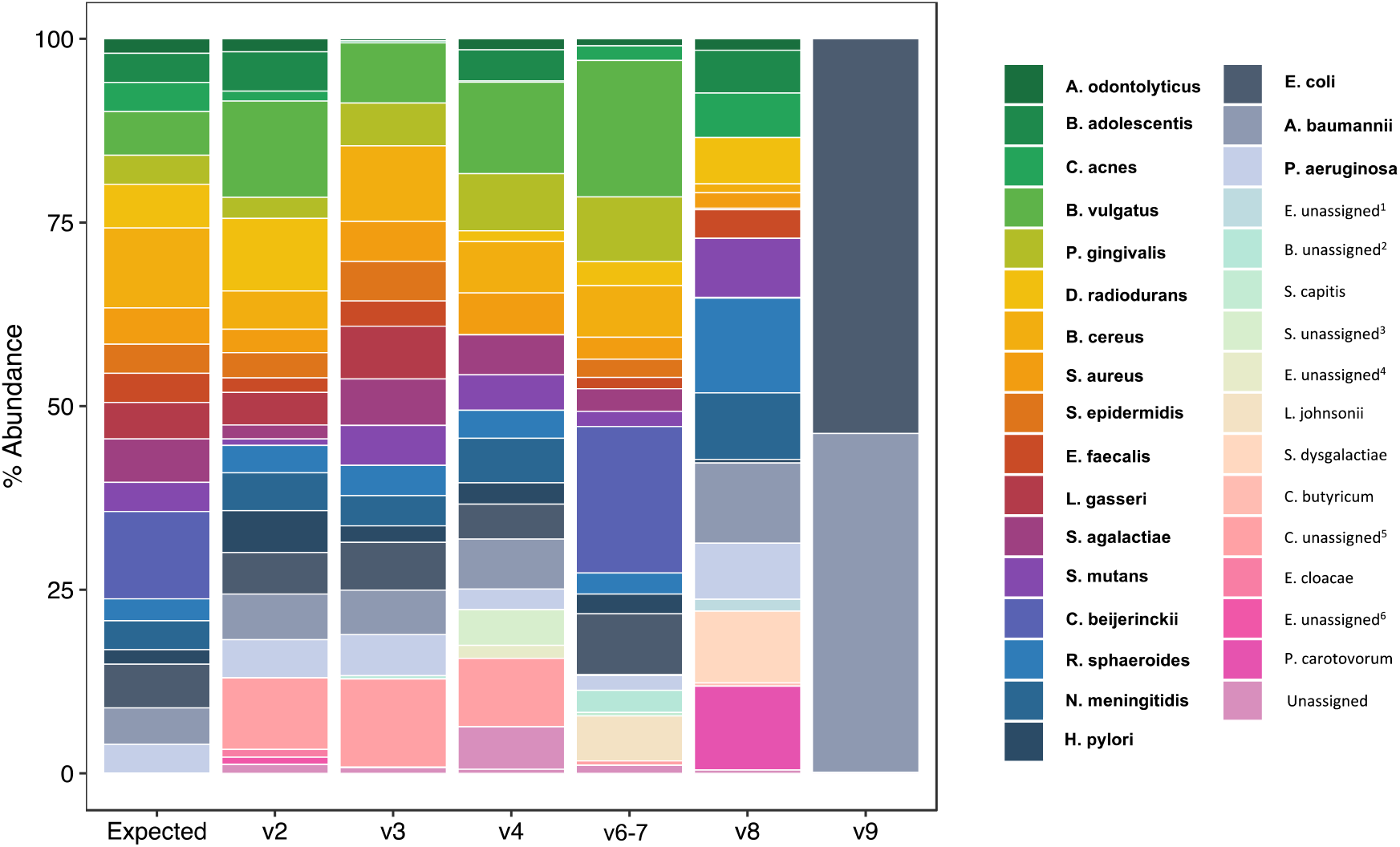
Species-level taxonomic barplots of ATCC 20 Strain Mix sequencing results by 16S rRNA hypervariable region. Bolded taxa are those present in the 20 strain mix. Enterobacteriaceae unassigned^1^, *Bifidobacterium* unassigned^2^, *Staphylococcus* unassigned^3^, *Enterococcus* unassigned^4^, *Clostridium sensu stricto 1* unassigned^5^, *Enterobacter* unassigned^6^.

**Table 3.**
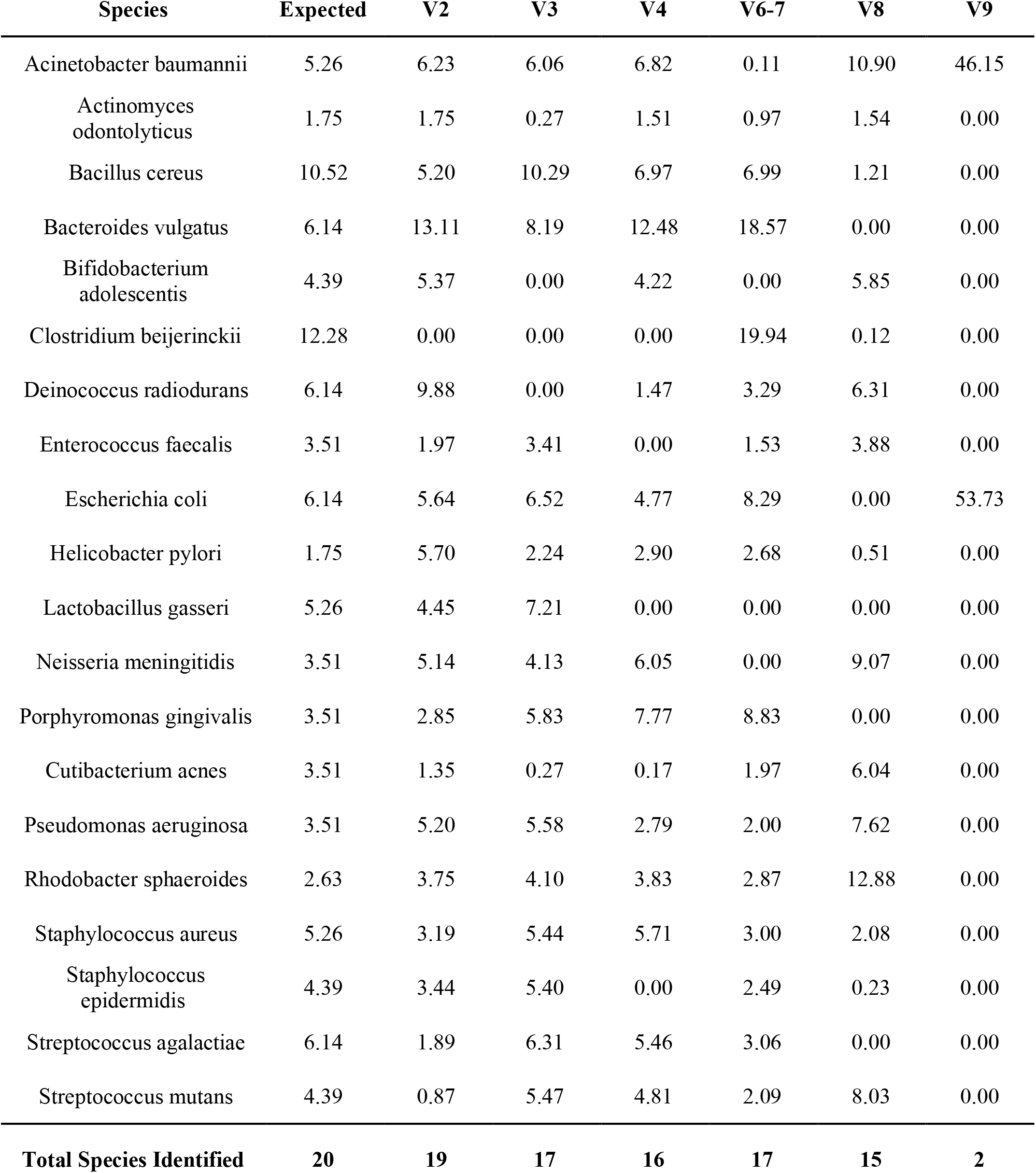
Observed species abundance denoted as percent of total.

Lastly, we performed a clustered heatmap analysis at the species level. The resulting heatmap demonstrated that technical replicates of the mock community sequences cluster by hypervariable region (Fig 4). The heatmap visually emphasizes the difference in taxonomic identification in V8 and particularly V9 compared to the other regions. It also highlights misclassifications and which regions were only able to classify taxa to the genus level.

**Figure 4.**
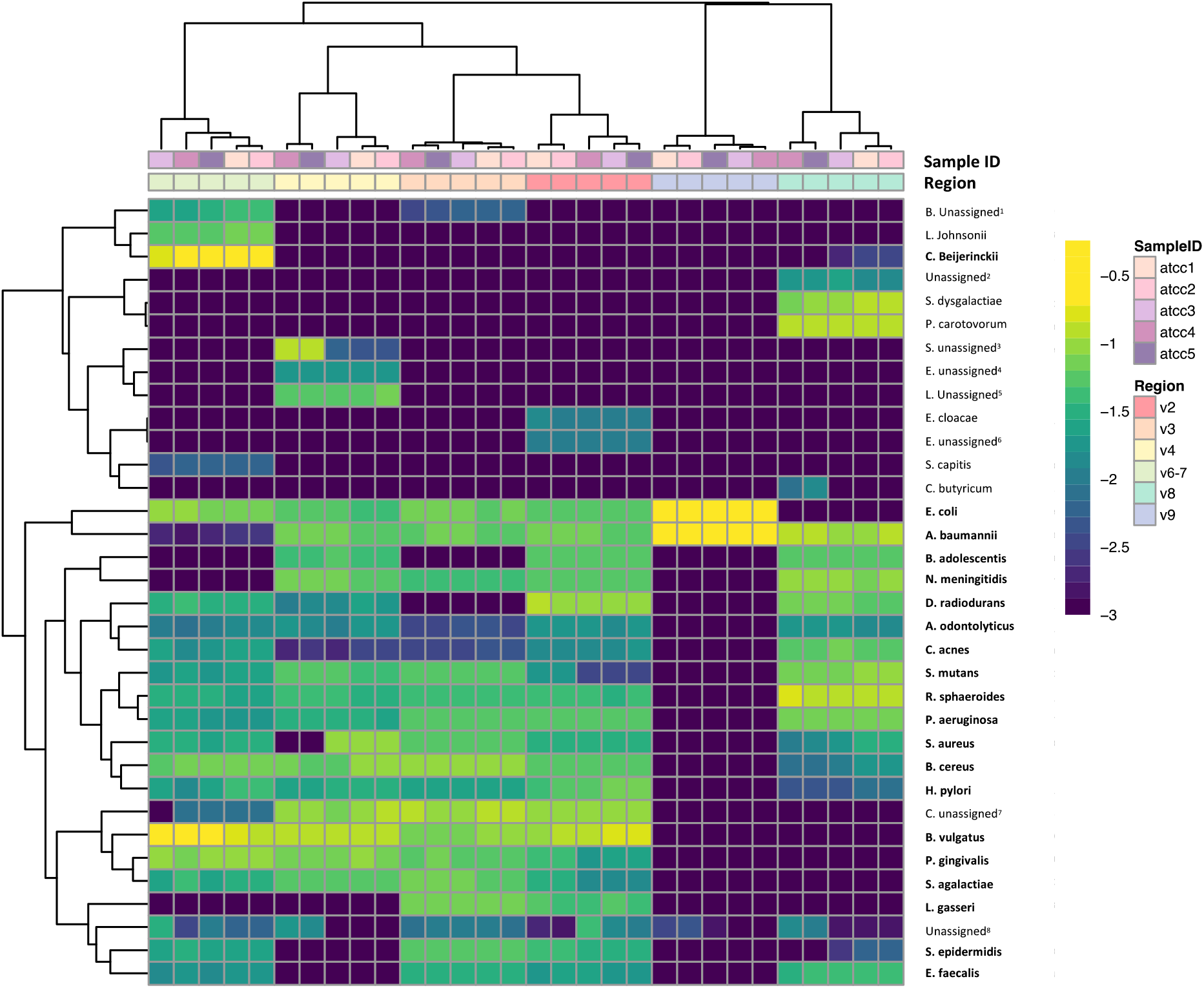
Species-level clustered heatmap of ATCC 20 Strain Even Mix. Bolded taxa are those present in the 20 strain mix. *Bifidobacterium* unassigned^1^, Enterobacteriaceae unassigned^2^, *Staphylococcus* unassigned^3^, *Enterococcus* unassigned^4^, *Lactobacillus* unassigned^5^, *Enterobacter* unassigned^6^, *Clostridium sensu stricto 1* unassigned^7^, Unassigned at every taxonomic level^8^.

### Clinical samples

We next sequenced and analyzed a set of six clinical samples in order to demonstrate the use of our pipeline on a clinical sample set. Samples consisted of duplicate rectal swabs from 3 participants. DNA was extracted immediately after collection from 1 rectal swab sample from each person (Fresh) and the other sample was frozen at −80 °C prior to DNA isolation (Frozen). Libraries were prepared in tandem, and all samples were sequenced on the same sequencing run. Sequencing results were processed as outlined above.

### Alpha diversity

We performed the same four alpha diversity metrics for the clinical cohort as for the mock community samples (Evenness, Shannon diversity, Observed OTUs, and Faith’s phylogenetic diversity). We used a GLM (see Methods) to incorporate data from hypervariable regions V2, V3, V4, and V6-7 in statistical analyses. We excluded data from the V8 and V9 regions due to the demonstrated poor performance of amplicons from these regions in identifying species in our *in silico* analysis and in the mock community (Table 2, Figs 1-4). There were no significant differences in alpha diversity between fresh and frozen samples by Shannon diversity, Faith’s phylogenetic diversity or Observed OTUs (Fig 5). Evenness was slightly increased in frozen samples across all hypervariable regions (adjusted GLM p = 0.015).

**Fig 5.**
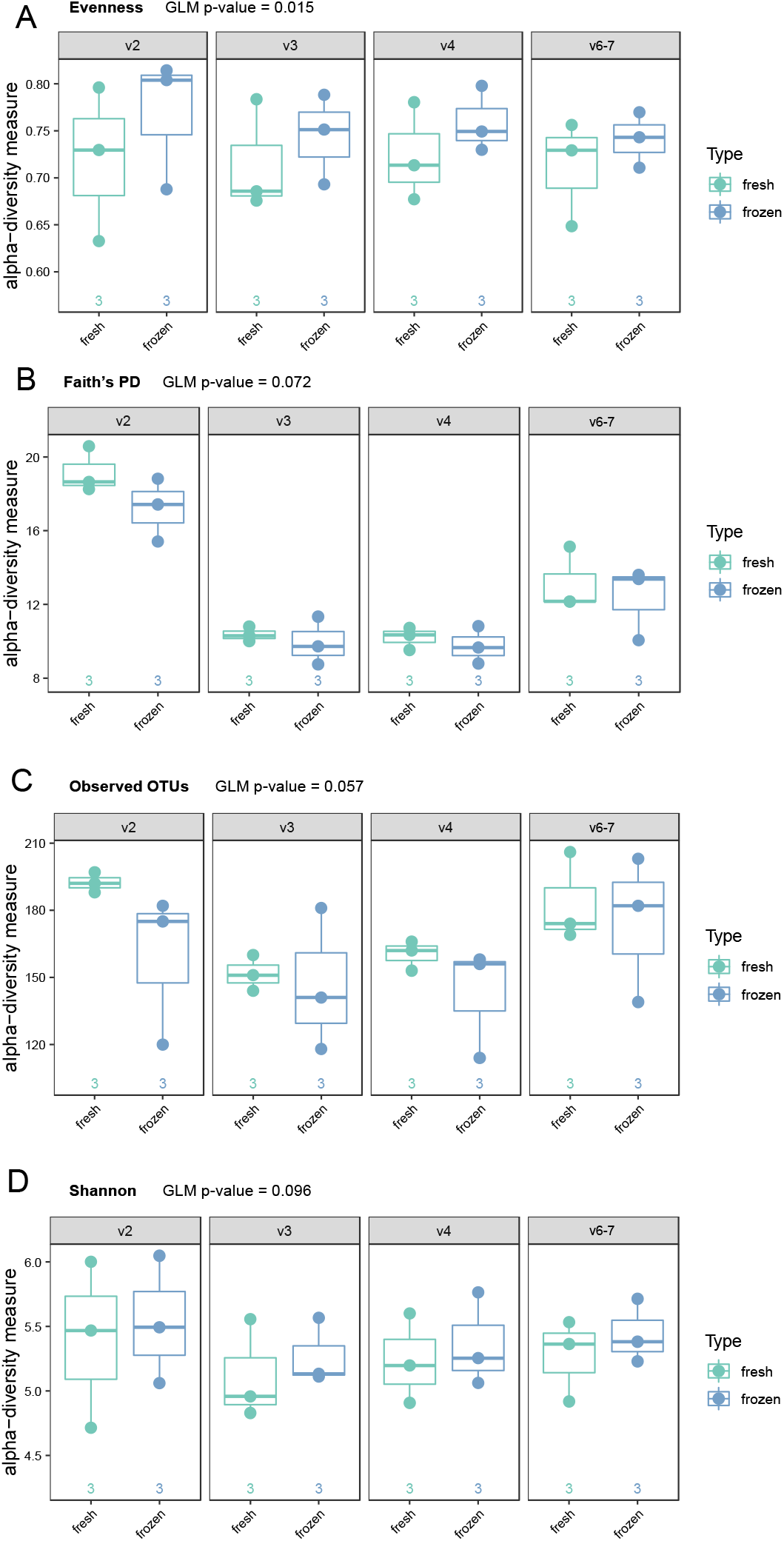
Alpha diversity analyses of six clinical samples by type (fresh or frozen) and hypervariable region. Each patient provided two swabs, one of which was frozen prior to DNA extraction. Evenness (A), Faith’s phylogenetic diversity (B), Observed Operational Taxonomic Units (OTUs) (C), Shannon diversity (D).

### Beta diversity

We aggregated taxonomic results and used them to create Bray-Curtis, Jaccard, Canberra, Euclidean, Gower, and Kulczynski distance matrices in order to perform combined beta diversity analysis across all hypervariable regions. As demonstrated by the Canberra PCoA plot in Fig 6, most variation in Beta diversity was due different individuals and V9 sequences. PERMANOVA analysis of results from each individual hypervariable region demonstrated that total composition does not differ by Fresh versus Frozen status after adjusting for individual person and region-to-region variation (S4 File).

**Fig 6.**
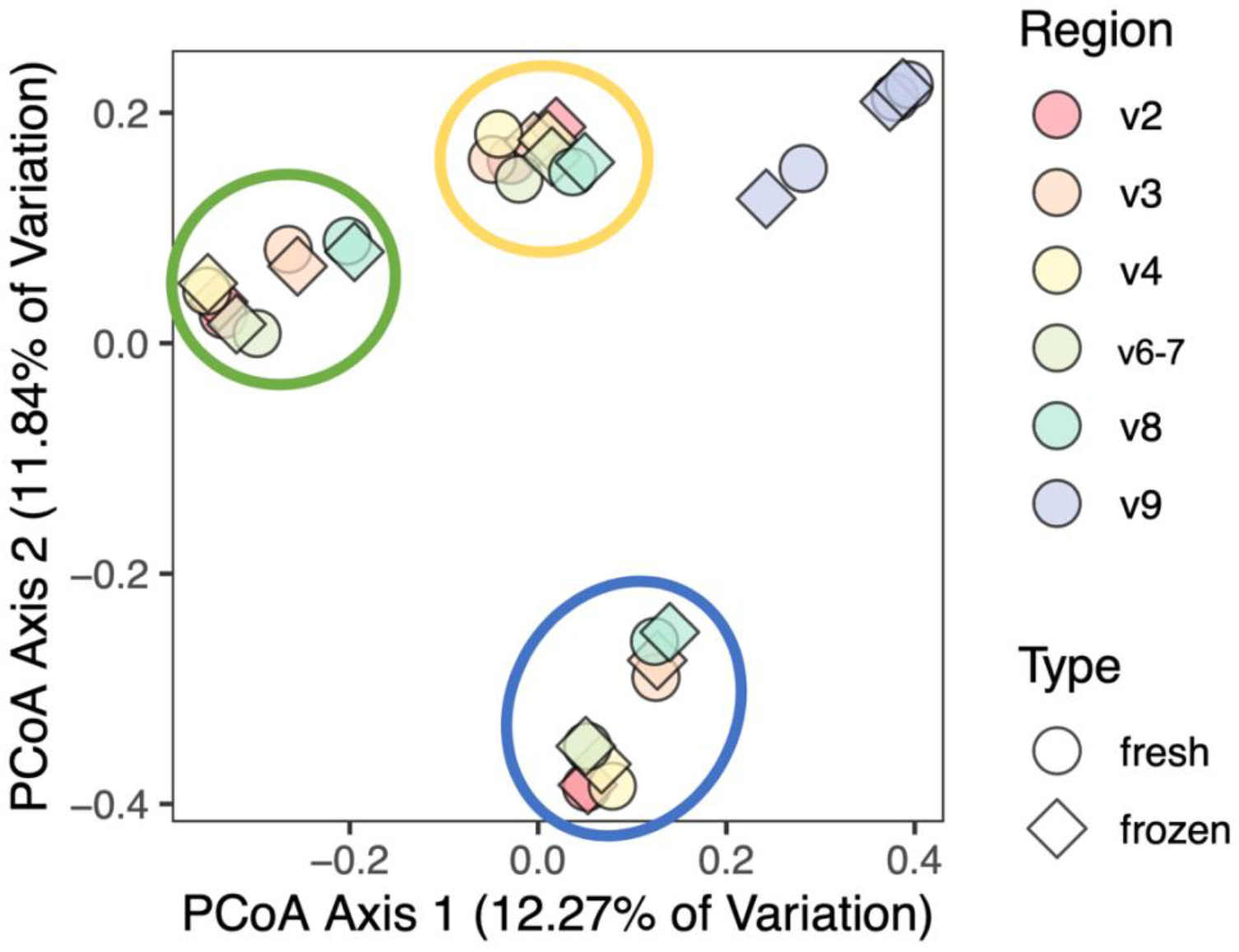
Principal coordinates analysis of clinical cohort using Canberra distance matrix. Samples and regions from the same person are circled, excluding V9. Results cluster by individual and by V9 region (not circled), but not by fresh versus frozen status.

### Taxonomic differential abundance

We systematically compared abundance of taxa between fresh and frozen samples at multiple levels (phylum, class, order, family, genus, species). We specifically looked at levels of Firmicutes, Bacteroidetes, and Faecalibacterium due to previous reports of differential abundance in fresh verses frozen samples (27, 28). Our results showed no significant differences in these taxa (Fig 7) or Firmicutes to Bacteroidetes ratios (Fig 8).

**Fig 7.**
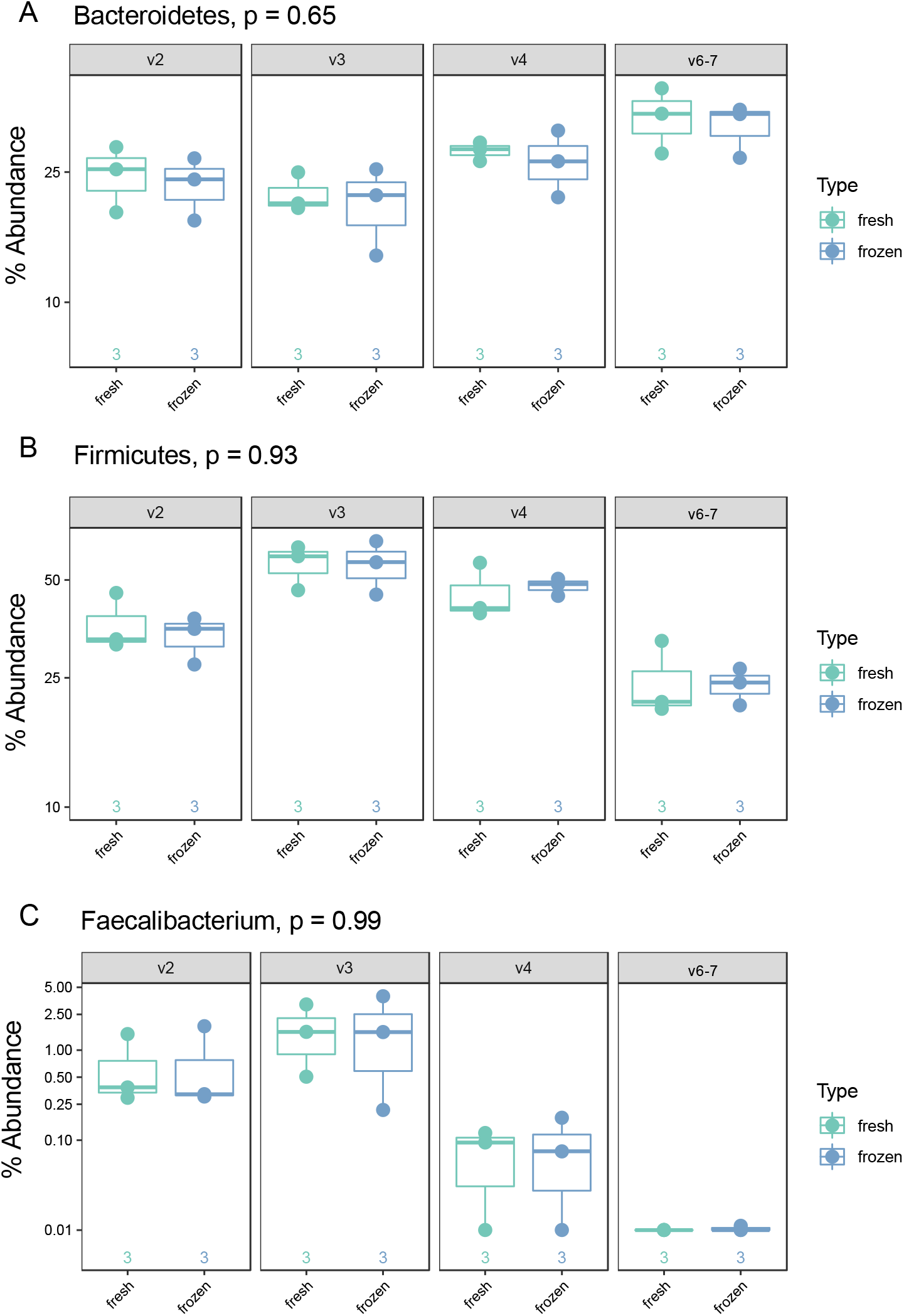
Percent abundance of Bacteroidetes, Firmicutes, and Faecalibacterium by sample type (fresh vs. frozen) and hypervariable region. P-value was calculated with a log-transformed GLM, and is false discovery rate-adjusted.

**Fig 8.**
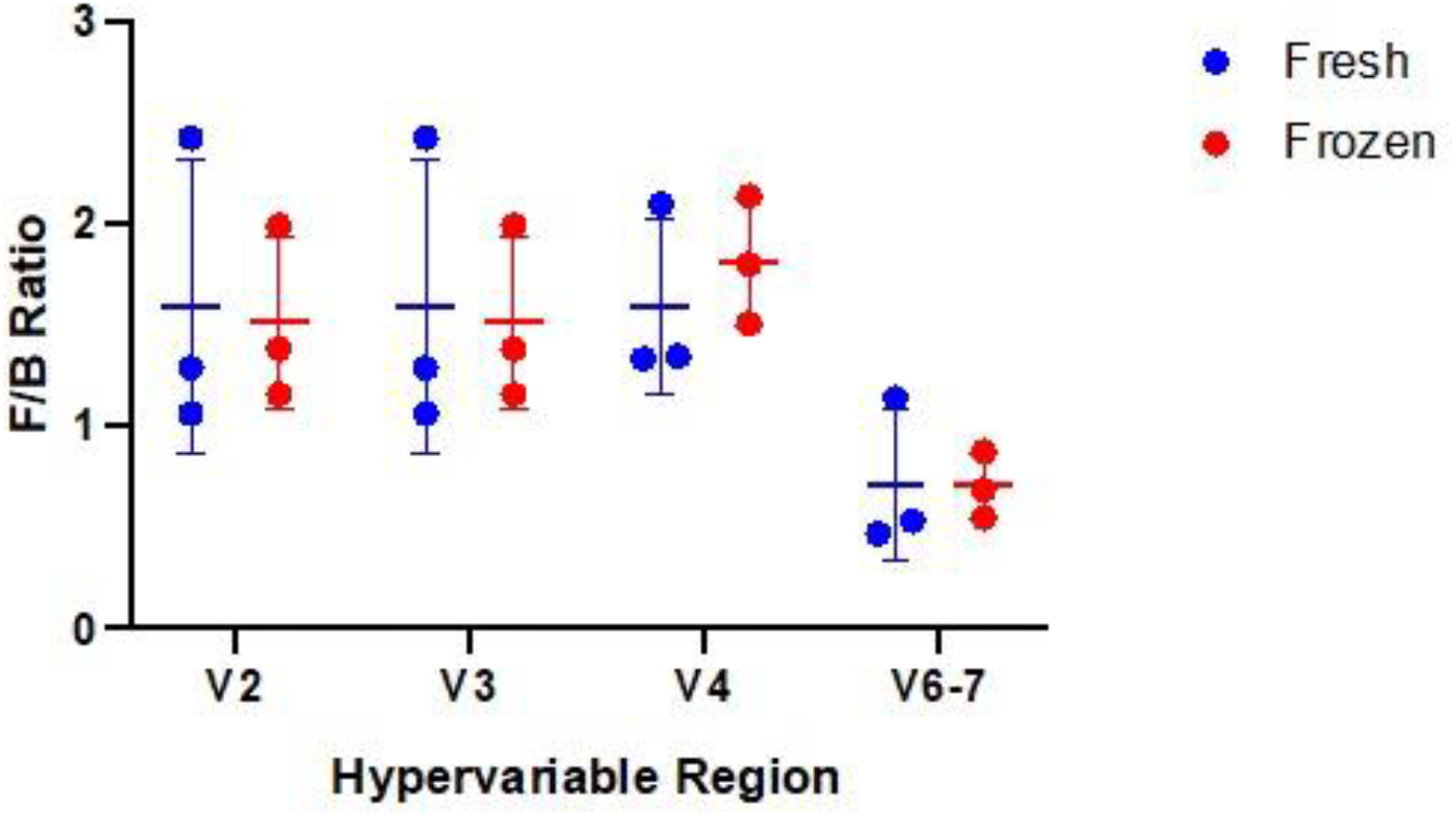
Comparison of Firmicutes to Bacteroidetes (F/B) ratio in fresh versus frozen samples by hypervariable region.

## Discussion

16S rRNA sequencing is a popular method for microbiome characterization due to multiple factors. The 16S rRNA gene encodes the small subunit of the prokaryotic ribosome and thus is present in the genome of all bacteria. The genetic sequence of 16S rRNA is composed of alternating conserved regions and nine hypervariable regions. PCR amplification using primers that target conserved regions and amplify across hypervariable regions allows amplification of DNA across a widespread taxonomic spectrum, and provides unique sequences that can be used for taxonomic classification at higher levels (e.g., family, genus, and species level). Next generation sequencing strategies are often limited to sequencing across only one or at most two of the nine hypervariable regions. The Ion 16S™ Metagenomics Kit provides the opportunity to prepare libraries containing sequences from seven of the nine hypervariable regions (V2, V3, V4, V6-7, V8, and V9). Herein we report results from sequencing a mock microbial community using the Ion 16S™ Metagenomics Kit and comparing results from different hypervariable regions. We also propose using a generalized linear multivariate model to incorporate sequencing data from multiple hypervariable regions for analyzing diversity and taxonomic abundances and demonstrate the use of this model in a small cohort of clinical samples.

The first step in our data analysis pipeline involved curating a taxonomic database. The Greengenes and SILVA databases are commonly used for taxonomic classification, however they are infrequently updated, inconsistently annotated, and contain redundant sequences (12, 13). Therefore, we refined the SILVA database by removing unclassified and distantly related sequences and formatting taxonomic annotations into a uniform layout to make sfanos db 4.0. Prior to using our database, we confirmed its accuracy by performing an *in silico* analysis using a human gut microbiome culture collection (14) where we ran known sequences through our classification scheme and recorded whether the sequences were classified correctly (true positive), incorrectly (false positive), or not identified (false negative). Encouragingly, our database correctly identified 85.05% of sequences to the genus level and 65.96% of sequences to the species level, which is an improvement compared to the rates previously published for the standard Greenegenes and SILVA databases (29). While our results are encouraging, it is important to keep in mind that nearly 44% of species-level calls are incorrect. Therefore, for both species-level and genus-level calls, we recommend that researchers consider reporting classification confidence scores along with results.

We next prepared and sequenced five technical replicates of DNA from a 20 strain mock microbial community and performed taxonomic classification. Even with our limited mock community dataset, we observed hypervariable region-based differences in alpha diversity. Most notably, taxa identified with V9 primers had significantly decreased alpha diversity compared to all other regions across all metrics. V8 results likewise had significantly decreased diversity across most metrics.

We performed six different beta diversity metrics (Bray-Curtis, Jaccard, Canberra, Euclidean, Gower, and Kulczynski) to evaluate differences between hypervariable regions. Distance matrices used in beta diversity analyses are generated from OTU tables, however the OTUs identified were not consistent among hypervariable regions. Therefore, in order to compare results between hypervariable region, we assembled distance matrices using taxonomic results. PCoA analyses demonstrated clustering primarily by hypervariable regions V2, V3, V4, and V6-7. Hypervariable regions V8 and V9 clustered separately from the other regions, again demonstrating the poor performance of amplicon sequencing of these regions in assessing the constituents of the mock community sample.

Consistent with previous reports (4–6, 8), we found that the taxonomic classification results from the mock community samples varied by hypervariable region. Primers targeting the V2, V3, and V6-7 regions identified nearly all the species present in the mock community (19/20, 17/20, and 17/20 respectively), V4 identified 16/20 species, V8 identified 15/20 species, and V9 identified only two (2/20) (Fig 3, Table 3). Generally, those regions which identified more species present in the mock community also had more evenly distributed observed taxa (i.e. there were no extreme over- or underestimated taxa which skewed the remaining percent abundances, such as in the case of V9).

The differences in the ability to identify taxa between hypervariable regions could be due to primer bias, database bias, or other biases in the analysis pipeline. Only 21 OTUs were assigned to the V9 region. Of those, two OTUs made up 99.78% of total V9 reads. Therefore, we deduce that the lack of diversity in the region is likely more related to primer bias than analysis or classification bias. Since V9 lacks sensitivity for many species, we opted to leave this region out of the generalized linear model we used on the clinical samples. V8 also tended to be less sensitive compared to V2, V3, V4, and V6-7, and did not cluster with the other regions on PCoA plots. Therefore, V8 was excluded from further analyses as well.

Researchers can circumvent the issue of choosing only one hypervariable region to analyze by sequencing multiple hypervariable regions in tandem. Since the sensitivity of each hypervariable region for identifying bacterial taxa varies, combining the results from multiple hypervariable regions for analyses may be misleading. Fuks et al. developed Short MUltiple Regions Framework (SMURF), which combines sequences from multiple PCR amplicons in order to provide one overall set of taxonomic profiling results (9). However, this method is computationally intensive and requires proprietary software. Therefore, to utilize information from multiple hypervariable regions at once and to strengthen confidence in the taxonomic abundance results, we incorporated a generalized linear model (GLM) into alpha diversity and taxonomic abundance analyses. We excluded V8 and V9 from GLM calculations because of the decreased sensitivity and specificity identified in our mock community analyses.

We demonstrated use of the GLM via analysis of a clinical cohort, where each participant donated two rectal swab samples, one of which was processed fresh and the other one frozen prior to DNA extraction. Alpha diversity analysis revealed increased Evenness in frozen samples compared to fresh samples. This trend was visualized in results from each individual hypervariable region and was strengthened in the GLM. There was no difference in Shannon’s diversity, Observed OTUs, and Faith’s phylogenetic diversity between fresh and frozen samples. Beta diversity analysis demonstrated clustering of samples by person irrespective of fresh versus frozen status or hypervariable region, with the exception of V9. PERMANOVA analysis confirmed that most of the variation in composition was due to individuals as opposed to storage type. An important limitation of our beta diversity analysis is that in order to compare results from all hypervariable regions in the same analysis, we had to use taxonomic classification as opposed to OTUs. This limits our beta diversity analysis to using only those reads that were assigned taxonomy.

Lastly, we compared taxonomic abundance at multiple levels between fresh and frozen samples and found no taxa at any level had significantly different abundance. This is unsurprising based on our small sample size, the fact that alpha and beta diversity were minimally different between sample type, and the fact that other studies show limited differences between fresh verses frozen samples (27, 28). However, Faecalibacterium results highlight the important point that not all regions are able to identify a taxon of interest: V6-7 fails to map any reads to this taxon despite its presence in the sample. Thus, even though the true composition of a clinical sample may be unknown, examining redundant data from multiple hypervariable regions may help elucidate the true microbial makeup of the sample.

In conclusion, we propose a method to overcome the issues of analyzing multiple amplicons covering multiple hypervariable regions at once. It is critical for the microbiome bioinformatics community to come to a consensus as to the proper way to analyze this type of data in order to maintain data quality, and to be able to compare results across different publications.

## Funding

This work was supported by The Assistant Secretary of Defense for Health Affairs Endorsed by the Department of Defense through the Prostate Cancer Research Program - Early Investigator Research Award under Award No. W81XWH-18-1-0545 (L.B.P.) and Prostate Cancer Challenge Award 16CHAL13 (K.S.S.). Opinions, interpretations, conclusions and recommendations are those of the author and are not necessarily endorsed by the Department of Defense.

## Acknowledgements

Thank you to the QIIME2 forum community for their help and discussions regarding analyzing multiple hypervariable regions of Ion Torrent data, especially Evan Bolyen, Nicholas Bokulich, Matthew Dillon, Justine Debelius, Colin Brislawn, Jennifer Barb, and Katherine Maki. We also appreciate the help and guidance from Kornel Schuebel, Hai Xu, and Jennifer Meyers of the JHU SKCCC Experimental and Computational Genomics Core; and Bradley Toms and Leonardo Varuzza at ThermoFisher.

## Supporting information

**S1 Fig. Other beta diversity metrics (Bray-Curtis, Jaccard, Euclidean, Gower, and Kulczynski), mock samples**

**S1 Table. Table of contaminants.**

**S1 File. In silico taxonomic validation results.**

**S2 File. Alpha diversity statistics.**

**S3 File. Filtered percent abundance.**

**S4 File. PERMANOVA analysis of fresh vs frozen clinical samples.**

## Notes

### Competing Interest Statement

JRW has financial and/or other relationship with Resphera Biosciences. This does not prevent us from sharing data and materials. There are no patents, products in development or marketed products associated with this research to declare.

## References

1. Quince C, Walker AW, Simpson JT, Loman NJ, Segata N. Shotgun metagenomics, from sampling to analysis. Nat Biotechnol. 2017;35(9):833–44.

2. Sanschagrin S, Yergeau E. Next-generation sequencing of 16S ribosomal RNA gene amplicons. J Vis Exp. 2014(90).

3. Ranjan R, Rani A, Metwally A, McGee HS, Perkins DL. Analysis of the microbiome: Advantages of whole genome shotgun versus 16S amplicon sequencing. Biochem Biophys Res Commun. 2016;469(4):967–77.

4. Cai L, Ye L, Tong AH, Lok S, Zhang T. Biased diversity metrics revealed by bacterial 16S pyrotags derived from different primer sets. PLoS One. 2013;8(1):e53649.

5. Barb JJ, Oler AJ, Kim HS, Chalmers N, Wallen GR, Cashion A, et al. Development of an Analysis Pipeline Characterizing Multiple Hypervariable Regions of 16S rRNA Using Mock Samples. PLoS One. 2016;11(2):e0148047.

6. Claesson MJ, Wang Q, O’Sullivan O, Greene-Diniz R, Cole JR, Ross RP, et al. Comparison of two next-generation sequencing technologies for resolving highly complex microbiota composition using tandem variable 16S rRNA gene regions. Nucleic Acids Res. 2010;38(22):e200.

7. Pinto AJ, Raskin L. PCR biases distort bacterial and archaeal community structure in pyrosequencing datasets. PLoS One. 2012;7(8):e43093.

8. Tremblay J, Singh K, Fern A, Kirton ES, He S, Woyke T, et al. Primer and platform effects on 16S rRNA tag sequencing. Front Microbiol. 2015;6:771.

9. Fuks G, Elgart M, Amir A, Zeisel A, Turnbaugh PJ, Soen Y, et al. Combining 16S rRNA gene variable regions enables high-resolution microbial community profiling. Microbiome. 2018;6(1):17.

10. Debelius JW, Robeson M, Hugerth LW, Boulund F, Ye W, Engstrand L. A comparison of approaches to scaffolding multiple regions along the 16S rRNA gene for improved resolution. bioRxiv. 2021:2021.03.23.436606.

11. Shrestha E, White JR, Yu SH, Kulac I, Ertunc O, De Marzo AM, et al. Profiling the Urinary Microbiome in Men with Positive versus Negative Biopsies for Prostate Cancer. J Urol. 2018;199(1):161–71.

12. Yilmaz P, Parfrey LW, Yarza P, Gerken J, Pruesse E, Quast C, et al. The SILVA and “All-species Living Tree Project (LTP)” taxonomic frameworks. Nucleic Acids Res. 2014;42(Database issue):D643–8.

13. McDonald D, Price MN, Goodrich J, Nawrocki EP, DeSantis TZ, Probst A, et al. An improved Greengenes taxonomy with explicit ranks for ecological and evolutionary analyses of bacteria and archaea. ISME J. 2012;6(3):610–8.

14. Forster SC, Kumar N, Anonye BO, Almeida A, Viciani E, Stares MD, et al. A human gut bacterial genome and culture collection for improved metagenomic analyses. Nat Biotechnol. 2019;37(2):186–92.

15. Bolyen E, Rideout JR, Dillon MR, Bokulich NA, Abnet CC, Al-Ghalith GA, et al. Reproducible, interactive, scalable and extensible microbiome data science using QIIME 2. Nat Biotechnol. 2019;37(8):852–7.

16. Callahan BJ, McMurdie PJ, Rosen MJ, Han AW, Johnson AJ, Holmes SP. DADA2: High-resolution sample inference from Illumina amplicon data. Nat Methods. 2016;13(7):581–3.

17. Good IJ. The population frequencies of species and the estimation of population parameters. Biometrika. 1953;40(3-4):237–64.

18. Bokulich NA, Dillon MR, Bolyen E, Kaehler BD, Huttley GA, Caporaso JG. q2-sample-classifier: machine-learning tools for microbiome classification and regression. J Open Res Softw. 2018;3(30).

19. Katoh K, Standley DM. MAFFT multiple sequence alignment software version 7: improvements in performance and usability. Mol Biol Evol. 2013;30(4):772–80.

20. Price MN, Dehal PS, Arkin AP. FastTree 2--approximately maximum-likelihood trees for large alignments. PLoS One. 2010;5(3):e9490.

21. Faith DP, Minchin PR, Belbin L. Compositional dissimilarity as a robust measure of ecological distance. Vegetatio. 1987;69(1-3):57–68.

22. Shannon CE. A mathematical theory of communication. ACM SIGMOBILE mobile computing and communications review. 2001;5(1):3–55.

23. Jaccard P. Nouvelles recherches sur la distribution florale. Bull Soc Vaud Sci Nat. 1908;44:223–70.

24. Sorensen TA. A method of establishing groups of equal amplitude in plant sociology based on similarity of species content and its application to analyses of the vegetation on Danish commons. Biol Skar. 1948;5:1–34.

25. Lozupone CA, Hamady M, Kelley ST, Knight R. Quantitative and qualitative beta diversity measures lead to different insights into factors that structure microbial communities. Appl Environ Microbiol. 2007;73(5):1576–85.

26. Lozupone C, Knight R. UniFrac: a new phylogenetic method for comparing microbial communities. Appl Environ Microbiol. 2005;71(12):8228–35.

27. Fouhy F, Deane J, Rea MC, O’Sullivan O, Ross RP, O’Callaghan G, et al. The effects of freezing on faecal microbiota as determined using MiSeq sequencing and culture-based investigations. PLoS One. 2015;10(3):e0119355.

28. Bahl MI, Bergstrom A, Licht TR. Freezing fecal samples prior to DNA extraction affects the Firmicutes to Bacteroidetes ratio determined by downstream quantitative PCR analysis. FEMS Microbiol Lett. 2012;329(2):193–7.

29. Agnihotry S, Sarangi AN, Aggarwal R. Construction & assessment of a unified curated reference database for improving the taxonomic classification of bacteria using 16S rRNA sequence data. Indian J Med Res. 2020;151(1):93–103.

