## Supplementary figures and images for "Incorporation of data from multiple hypervariable regions when analyzing bacterial 16S rRNA sequencing data"

### Supplementary Figure 1

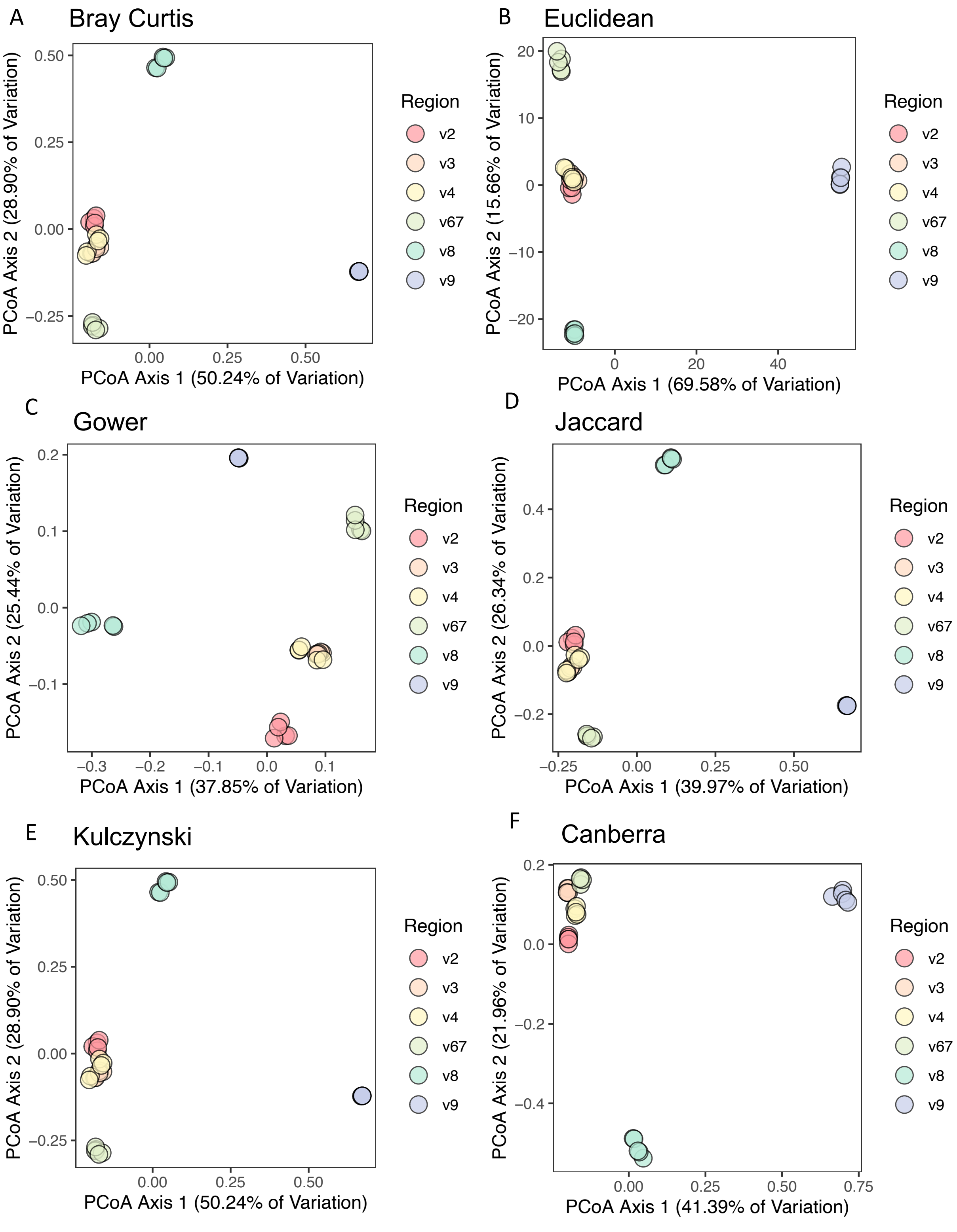
