## Supplementary Table 1 for "Incorporation of data from multiple hypervariable regions when analyzing bacterial 16S rRNA sequencing data"

**S1 Table. List of contaminants.**

| Contaminant | Region Detected | Reason for Filtering |
|---|---|---|
| k__Bacteria; p__Actinobacteria; c__Actinobacteria; o__Actinomycetales; f__Actinomycetaceae; g__Varibaculum; s__unassigned | V67 | Only detected in 1 replicate; less than 0.1% abundance |
| k__Bacteria; p__Actinobacteria; c__Actinobacteria; o__Corynebacteriales; f__Corynebacteriaceae; g__Corynebacterium_1; s__Corynebacterium_amycolatum | V67 | Only detected in 1 replicate; less than 0.1% abundance |
| k__Bacteria; p__Actinobacteria; c__Actinobacteria; o__Micrococcales; f__Microbacteriaceae; g__Microbacterium; s__unassigned | V3, V67 | Only detected in 1 replicate; less than 0.1% abundance |
| k__Bacteria; p__Bacteroidetes; c__Bacteroidia; o__Bacteroidales; f__Bacteroidaceae; g__Bacteroides; s__Bacteroides_dorei | V8 | Less than 0.1% abundance |
| k__Bacteria; p__Bacteroidetes; c__Bacteroidia; o__Bacteroidales; f__Bacteroidaceae; g__Bacteroides; s__Bacteroides_faecis | V3 | Only detected in 1 replicate; less than 0.1% abundance |
| k__Bacteria; p__Bacteroidetes; c__Bacteroidia; o__Bacteroidales; f__Bacteroidaceae; g__Bacteroides; s__Bacteroides_fragilis | V3 | Only detected in 1 replicate; less than 0.1% abundance |
| k__Bacteria; p__Bacteroidetes; c__Bacteroidia; o__Bacteroidales; f__Bacteroidaceae; g__Bacteroides; s__Bacteroides_massiliensis | V67 | Only detected in 1 replicate; less than 0.1% abundance |
| k__Bacteria; p__Bacteroidetes; c__Bacteroidia; o__Bacteroidales; f__Bacteroidaceae; g__Bacteroides; s__Bacteroides_thetaiotaomicron | V67 | Only detected in 1 replicate; less than 0.1% abundance |
| k__Bacteria; p__Bacteroidetes; c__Bacteroidia; o__Bacteroidales; f__Porphyromonadaceae; g__Parabacteroides; s__Parabacteroides_merdae | V3, V67 | Less than 0.1% abundance |
| k__Bacteria; p__Firmicutes; c__Bacilli; o__Bacillales; f__Bacillaceae; g__Bacillus; s__unassigned | V9 | Only detected in 1 replicate |
| k__Bacteria; p__Firmicutes; c__Bacilli; o__Bacillales; f__Family_XI; g__Gemella; s__Gemella_morbillorum | V3 | Only detected in 1 replicate; less than 0.1% abundance |
| k__Bacteria; p__Firmicutes; c__Clostridia; o__Clostridiales; f__Family_XI; g__Anaerococcus; s__Anaerococcus_vaginalis | V3 | Only detected in 1 replicate; less than 0.1% abundance |
| k__Bacteria; p__Firmicutes; c__Clostridia; o__Clostridiales; f__Family_XI; g__Ezakiella; s__Sporobacterium_sp | V3, V67 | Only detected in 1 replicate; less than 0.1% abundance |
| k__Bacteria; p__Firmicutes; c__Clostridia; o__Clostridiales; f__Family_XI; g__Finegoldia; s__Finegoldia_magna | V3 | Only detected in 1 replicate; less than 0.1% abundance |
| k__Bacteria; p__Firmicutes; c__Clostridia; o__Clostridiales; f__Family_XI; g__Peptoniphilus; s__Candidatus_Peptoniphilus_massiliensis | V3, V67 | Only detected in 1 replicate; less than 0.1% abundance |
| k__Bacteria; p__Firmicutes; c__Clostridia; o__Clostridiales; f__Lachnospiraceae; g__Anaerostipes; s__Anaerostipes_hadrus | V67 | Only detected in 1 replicate |
| k__Bacteria; p__Firmicutes; c__Clostridia; o__Clostridiales; f__Lachnospiraceae; g__Blautia; s__unassigned | V3 | Only detected in 1 replicate |
| k__Bacteria; p__Firmicutes; c__Clostridia; o__Clostridiales; f__Lachnospiraceae; g__Lachnoclostridium; s__unassigned | V3 | Only detected in 1 replicate; less than 0.1% abundance |
| k__Bacteria; p__Firmicutes; c__Clostridia; o__Clostridiales; f__Lachnospiraceae; g__unassigned; s__unassigned | V4 | Only detected in 1 replicate |
| k__Bacteria; p__Firmicutes; c__Clostridia; o__Clostridiales; f__Ruminococcaceae; g__Ruminococcus_1; s__Ruminococcus_bicirculans | V8 | Only detected in 1 replicate |
| k__Bacteria; p__Proteobacteria; c__Alphaproteobacteria; o__Rhizobiales; f__Methylobacteriaceae; g__Methylobacterium; s__unassigned | V3 | Only detected in 1 replicate; less than 0.1% abundance |
| k__Bacteria; p__Proteobacteria; c__Betaproteobacteria; o__Burkholderiales; f__Burkholderiaceae; g__Cupriavidus; s__Cupriavidus_metallidurans | V2, V4 | Less than 0.1% abundance |
| k__Bacteria; p__Proteobacteria; c__Deltaproteobacteria; o__Desulfovibrionales; f__Desulfovibrionaceae; g__Bilophila; s__Bilophila_wadsworthia | V9 | Only detected in 1 replicate |
